# Interhemispheric connectivity endures across species: An allometric exposé on the corpus callosum

**DOI:** 10.1101/2021.01.10.426155

**Authors:** Ben Cipollini, Garrison W. Cottrell

## Abstract

Rilling & Insel have argued that in primates, bigger brains have proportionally fewer anatomical interhemispheric connections, leading to reduced functional connectivity between the hemispheres (1). They based this on a comparison between surface areas of the corpus callosum and cortex rather than estimating connection counts, while leaving out other quantities also dependent on brain size such as callosal fiber density, neuron density, and number of functional areas.

We use data from the literature to directly estimate connection counts. First, we estimate callosal fiber density as a function of brain size. We validate this by comparing out-of-sample human data to our function’s estimate. We then mine the literature to obtain function estimates for all other quantities, and use them to estimate intra- and interhemispheric white matter connection counts as a function of brain size.

The results show a much larger decrease in the scaling of interhemispheric to intrahemispheric connections than previously estimated. However, we hypothesize that raw connection counts are the wrong quantity to be estimating when considering functional connectivity. Instead, we hypothesize that functional connectivity is related to connection counts relative to the number of cortical areas.

Accordingly, we estimate inter-area connection counts for intra- and interhemispheric connectivity and find no difference in how they scale with brain size. We find that, on average, an interhemispheric inter-area connection contains 3-8x more connections than an intrahemispheric inter-area connection, regardless of brain size. In doing so, we find that the fiber count of the human corpus callosum has been underestimated by 20%.

**Significance Statement:** There are arguments in the literature that larger brains have proportionally fewer interhemispheric connections. We find that the decrease is even larger than previously estimated. However, we argue that this quantity is the wrong thing to measure: Rather, we should measure functional connectivity between cortical areas. We show that the ratio of interhemispheric and intrahemispheric connectivity between cortical areas is constant across mammalian species. These findings are consistent with a growing literature that suggest interhemispheric connectivity is special across all primate species.

---

**L**ateralization touches on virtually every function that we believe humans are unique good at, including language, fine motor skills, spatial cognition, and social perception. One influential idea in the literature states that lateralization in humans is a consequence of greater independence between our cerebral hemispheres, driven by architectural (2) and physiological (3) differences purely due to increased brain size.

There are a number of reasons to doubt the hypothesis that large brains cause reduced interhemispheric communication, which in turn causes lateralization. First, interhemispheric connections appear to be special: they show greater functional correlation than intrahemispheric connections in humans (4), and this correlation does not appear to be a function distance (5) as it is with intrahemispheric connections (5, 6). This suggests that these connections are so important that they are maintained despite the large volumetric cost of their long length (7). Second, lateralization and interhemispheric communication are positively correlated in human development (8–10) and in neural network models of hemispheric interaction (11), suggesting a mutually reinforcing relationship between lateralization and interhemispheric communication rather than an antagonistic one. Finally, there is no direct evidence of decreased communication between the hemispheres when directly compared though oscillations (12, 13), in modeling of lateralization (11) or conduction delay (14).

Here, we re-analyze one of the two datasets used in support of the hypothesis that humans have greater hemispheric independence due to our large brain size. Rilling & Insel (2) used MRI to scan brains from 11 different primate species. To examine how total interhemispheric connectivity scales with respect to total white matter connectivity, they estimated the mid-sagittal callosal surface area (as a proxy for interhemispheric connectivity) and grey matter surface area (as a proxy for neuron count, and therefore total white matter connectivity). They found that, while callosal surface area increases with brain size, it increases at a slower rate than the grey matter surface area. They concluded that total interhemispheric connectivity is selectively and specifically reduced with increasing brain size, leading to decreased communication across the hemispheres with increasing brain size.

There are two potential weaknesses in the inferences made from these data. First, the relationship between these surface area measurements and actual connection counts depends on other quantities known to scale with brain size, such as neuron density in the grey matter (15) or axon packing density in the white matter (16–19). The scaling relationship of actual connections is therefore unlikely to follow that of the surface areas computed by Rilling & Insel. Second, we question whether comparison of total intrahemispheric and interhemispheric white matter connectivity reflects any meaningful functional measure of cortical architecture. Instead, we suggest that the scaling of fiber counts that interconnect cortical areas–inter-area fiber counts–is a better anatomical correlate of functional communication, and one that is more commonly measured in the literature (6, 20, 21). There are two potential weaknesses in the inferences made from these data. First, the relationship between these surface area measurements and actual connection counts depends on other quantities known to scale with brain size, such as neuron density in the grey matter (15) or axon packing density in the white matter (16–19). The scaling relationship of actual connections is therefore unlikely to follow that of the surface areas computed by Rilling & Insel. Second, we question whether comparison of total intrahemispheric and interhemispheric white matter connectivity reflects any meaningful functional measure of cortical architecture. Instead, we suggest that the scaling of fiber counts that interconnect cortical areas–inter-area fiber counts–is a better anatomical correlate of functional communication, and one that is more commonly measured in the literature (6, 20, 21).

In order to address these issues, we formulated three simple equations for estimating interhemispheric and total white matter connection counts from Rilling & Insel’s data (Equations 1-3). We collect the necessary data from the literature (using primate data almost exclusively; see (22, 23)), normalize them, and compute connection count estimates.

We find that Rilling & Insel were correct in their conclusion that total interhemispheric white matter connectivity decreases faster than total intrahemispheric connectivity; in fact, we find that the rate of decrease is much greater than the value they reported. This decrease is so strong that, given the literature demonstrating the importance of inter-hemispheric connections, it made us doubt that total connection counts is a good estimator of changes in functional communication.

We propose that comparing connection counts in fiber bundles between *functional areas* is a more relevant measure of communication. When we compare the scaling of connection counts in intra- vs. interhemispheric area-area fiber bundles, we find that inter- and intrahemispheric functional connectivity scale virtually identically. We also find evidence that interhemispheric inter-area connections contain more fibers than the average intrahemispheric inter-area connections, suggesting that they play a special role not only in humans (5), but in all primate species, regardless of brain size.

These analyses, based on the best available data, contradict the conclusions of Rilling & Insel, and instead support the emerging idea that callosal connections are of great importance in primates. While further testing of our hypothesis is warranted, it is clear that it provides a strong alternative to the view that humans have reduced interhemispheric communication, and that reduced interhemispheric communication leads to lateralization. We also hope that our review of the current available data helps locate important gaps in the empirical literature that need to be filled to come to firmer conclusions. Finally, we also hope that our careful identification of confounds due to age help foster a more careful consideration of how data are being used across studies.

## Comparing estimates of total white matter connectivity

The quantities estimated by Rilling & Insel, callosal and grey matter surface areas, can be related to total white matter connectivity as follows:

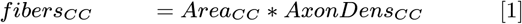

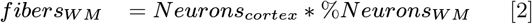

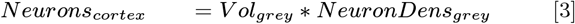

where *fibersCC* is the total number of white matter connections crossing the corpus callosum, *AreaCC* is the mid-sagittal surface area of the corpus callosum, *AxonDensCC* is the number of axons per unit area of the corpus callosum, *fibersW M* is the total number of white matter connections in the brain, *Neuronscortex* is the number of neurons in cortex.

Intuitively, Equation 1 says that the total number of callosal connections depends on the surface area of the callosum multiplied by the density (per unit area) of fibers in the callosum. Equations 2 and 3 say that the total number of white matter connections, regardless whether they interconnect cortex within or across hemispheres, depends on the total number of neurons (grey matter volume times the density of neurons per unit volume) and the percentage of neurons that project into the white matter^∗^.

All of the quantities needed for these computations have been measured across species, and all but callosal axon density have been characterized using allometry. Here, we use the best available allometric estimates from the literature, along with a novel estimate of callosal axon density, to estimate total white matter connections across the corpus callosum and cortex as a whole.

In order to estimate our allometric estimate of callosal axon density, we used data from a sample of six small-brained species (24). Wang et al. (2008) processed their samples with the best imaging and fixation procedures available, and used preparations at similar sample ages, hence these are the best available data. These data show the same kind of log-normal / “fat tail” distributions described in many places (e.g. (25, 26)). We used these data to compute an allometric relationship between callosal axon density and brain size.

Because these data are for smaller-brained species than the human species, we use corrected data from the best available human data (27) to see how callosal axon density for humans compares to our allometric estimate. In order to do so, we apply published corrections for tissue fixation and imaging, and derive a novel age correction based on high-quality monkey data (28). In doing so, we point out that most frequently quoted fiber count of the human corpus callosum (200 million) is confounded by age; our estimates suggest an age-corrected value of 240 million fibers − 20% larger than the commonly reported figure.

### Allometric regression of animal callosal axon density

Most research on the corpus callosum has focused on mid-sagittal area size, looking for correlations with brain size, intelligence, asymmetry, handedness, gender, and diseases (29). Far fewer resources have been put into examining the microstructure of the corpus callosum using microscopic techniques. These studies have pointed out the importance of tissue preservation and the use of electron microscopy, as the corpus callosum contains many thin, unmyelinated fibers (27) that disintegrate without careful tissue preservation and cannot reliably be imaged with a light microscope. Unfortunately, only a handful of studies use the electron microscope at all (16–18, 24, 30), and neither of the two studies on human tissue (27, 31) have preserved tissue as well as is done for non-human samples (32) due to differences in time since death and other procedural differences. In addition, human samples were obtained from much older individuals than is typical in animal studies, both in number of years and percentage of species-average life span.

## Materials & Methods

Wang et al. (2008) share the best data for conducting allometric regression, due to their carefully-reported procedures (24). They used electron microscopy to examine six small-brained species, carefully reported tissue processing and imaging procedures, and reported brain sizes (24). Though Wang’s data mixes rodent and primate data, the best available data suggests that axon caliber data is similar between the genera (22), and therefore we believe we’re on safe ground to use it.

Unfortunately, as with each dataset we wished to use, the raw data were unavailable for a variety of reasons. The data we used for our analyses were estimated from the figures in the pdf of the publication, using digital processing to estimate values from the published scatter plots and histograms.

Our procedure for extracting data from the pdf is as follows: Density data are plotted as a function of brain diameter on Wang et al. Figure 1e, with individual samples and species averages for all six species. Average brain weights for the 6 species were published in Wang et al. Table S1. On a 27” screen (iMac), we zoomed into the figure to maximize the limiting dimension (usually vertical), then captured a screen shot of the figure. We used the mouse to mark each X- and Y-axis tick-mark, the X- and Y-axis locations, and the centroid of each data point (see Figure S1 for an example). Each data point’s centroid was computed as the mean pixel within the line; when the line contained an even number of pixels, we chose the left-most (X-axis) or bottom-most (Y-axis) of the middle two pixels.

**Fig. 1.**
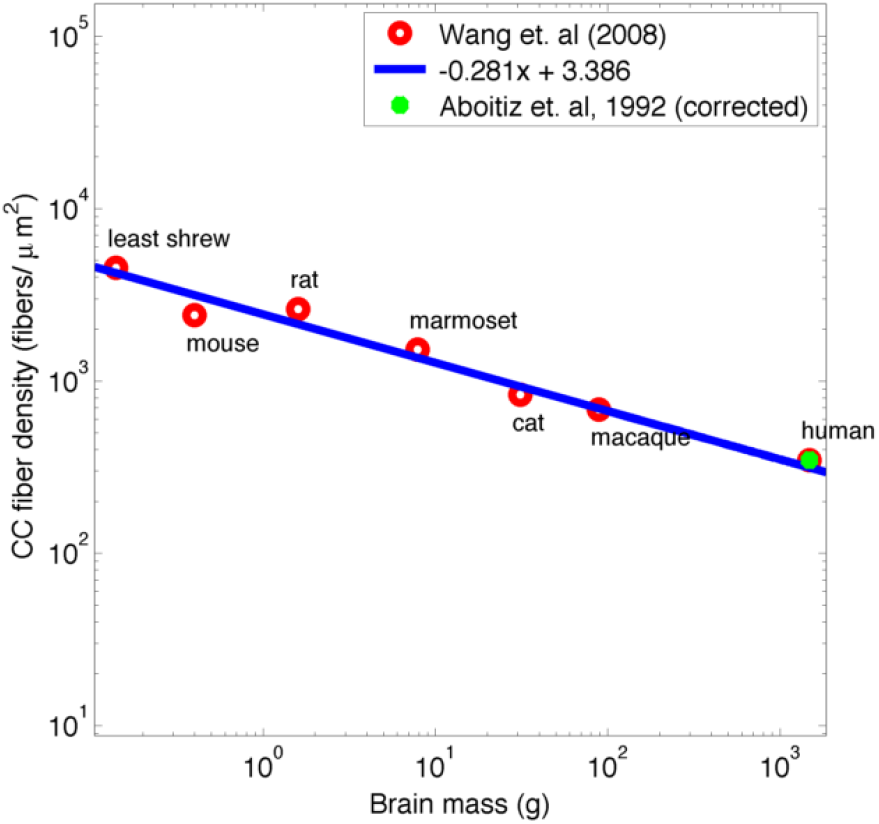
Allometric regression of callosal fiber density (blue line) using data from Wang et al. (red circles), on log-log axes, shows that callosal axon density decreases with brain size (slope: −0.281). The green data point indicates corrected value for Aboitiz et al. human data, and was not used to fit the regression line.

We developed MATLAB (R2010a) scripts read each annotated image and parse out values. Data estimates, as well as parsing code, are available online^†^; the manually annotated figures are not publicly available due to copyrights on the original figures. We validated the extracted values by estimating summary statistics or regression coefficients from them, and then compared these estimates to reported values in the publications. The error on the estimated summary statistics is generally less than 2%; see Table S1 for all error estimates.

We computed an allometric regression of corpus callosum axon density by regressing the *log*10 estimates of brain weight and corpus callosum density using reduced major axis regression (33). Reduced major axis regression allows for measurement error in both the X- and Y-axis values; ordinary least-squares regression assumes noiseless values for the X-axis.

## Results and Discussion

Using data from Wang et al., we find that callosal fiber density decreases with an exponent of *−* 0.281 with respect to brain mass (*r*^2^ = 0.909; see Figure 1). As expected, corpus callosum density scales negatively with brain size, consistent with data showing greater myelination and thicker axons in species with larger brains (19, 24, 34). This is the first time a quantitative regression of callosal axon density has been reported. Previous regressions either regress callosal area (25) or a subset of total fibers, such as regressing only on the largest fibers in the sample (19, 35).

### Comparing callosal estimates to age-corrected human data

Only small brains were used to measure callosal fiber density; the largest brain used by Wang et al. was the rhesus macaque monkey brain (89 grams), 15x smaller than the average human brain (1350 grams). This could make the regression inappropriate for larger-brained species.

In order to compare the allometric regression to human data, we searched for human electron microscopic data in the literature. Though a few studies have been published using a light microscope (27, 36–39), we only found data on one small human sample that has been imaged using an electron microscope (27)^‡^. Aboitiz et al. reported callosal axon density from a light microscope (371.7 fibers/*µ*m^2^), and estimated from a single sample that 20% additional small-caliber axons (*<* 1*µ*m) were found using an electron microscope. Aboitiz et al. used this estimate to correct their light-microscope density estimates; we use this correction factor here.

To compare these data to the Wang et al. data, we additionally had to correct for differences in tissue fixation procedures and sample age. Aboitiz et al. reported 35% tissue shrinkage (vs. 0% for Wang et al.) and a mean sample age of 45 years old (vs. “young adult” samples for Wang et al.). A multiplicative correction for shrinkage was reported, but differences in sample age have not been noted previously, nor any correction procedure suggested. The literature indicates that younger samples should have greater densities. Callosal density decreases in early post-natal development as fibers are pruned, axon diameters increase, and myelination occurs (16, 17). We analyzed adult data in macaques obtained from LaMantia & Rakic (17) and found a decrease in axon density that continues beyond sexual maturity (see Figure 2).

**Fig. 2.**
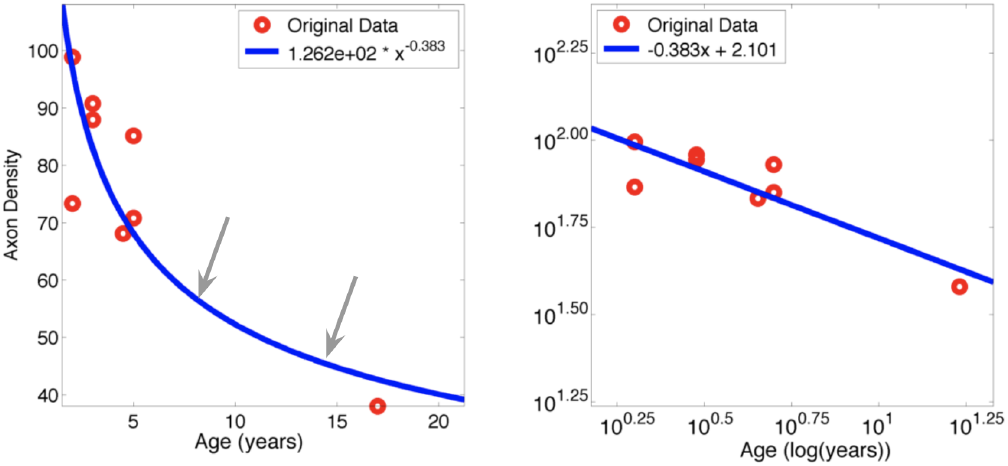
Left: changes in axon density for macaque adults, plotted on linear axes and with the computed allometric regression (blue line). The average adult macaque age (left arrow; 5.2 years old) and the human sample age (45 years old) mapped onto macaque lifespan (right arrow; 14.8 years old). See the text for detailed methods. Right: Same data and regression, with log-log axes. Data from LaMantia & Rakic (18).

We expected the human data to match or exceed the estimate from our allometric regression. A number of data indicate that humans have more small-diameter fibers than would be expected based on brain size alone. First, corpus callosum size in humans correlates with small diameter fiber count (27), but not in the smaller-brained macaque monkey (19), suggesting a larger proportion of small diameter axons in the human brain. Second, small diameter fibers are associated with association cortices such as temporal and prefrontal cortices (40), and comparative studies suggest that the major areas for human cortical expansion is in association cortices (41). Finally, Caminiti et al. (34) compared axon diameter distributions (ADD) between macaque, chimpanzee, and human brains and found that while the chimpanzee ADD compensates for the larger brain size, the human one does not.

## Materials & Methods

We apply three corrections to the light correction, as the density decrease from the younger samples is microscope human data from Aboitiz et al. (27). Two are multiplicative factors reported in the paper: 20% additional fibers found using electron microscopy over light microscopy and 35% tissue shrinkage, the latter of which means density is overestimated by 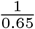. The third is a novel correction for differences across samples due to age, which we describe below.

We use allometric scaling to correct for differences in sample age by deriving an allometric curve from macaque electron microscopy data (17). We then followed the methods of Finlay et. al for cross-species remapping (42) to map the human sample age onto the macaque lifespan (see Figure 2), by using data on key maturational ages in each species to map data across species. We then use the points on the allometric curve for mean macaque sample age and “macaque-mapped” mean human age to compute a multiplicative factor for the expected difference between the samples, due to the differences in life stage. We then correct for this life stage confound using this newly computed multiplicative correction.

Following Finlay et al’.s procedure to map the human sample age onto the macaque lifespan, we obtained values for sexual maturity and average age of death for the two species (macaque: sexual maturity at 4 years old (yo), lifespan 25 yo, human: sexual maturity at 15 yo, lifespan 73 yo)(43). We computed what percentage of the human lifespan between sexual maturity and death the human data sample age (45 years) corresponds to by using the following equation: ^1^ *pct* = (*age*_*human*_ *−* 15)*/*(73 *−* 15). We then mapped that percentage onto the macaque lifespan between sexual maturity and death using the following equation: *age* = 4 +*pct∗* (25 *−* 4).

We also compute a prediction for human corpus callosum density, based on the average human brain weight and the allometric estimate from the Wang et al. data. We use average human brain weight of 1350 grams (43).

## Results

As shown in Figure 2b, the regression on age gives an equation *y* = 6.1 * *x −* 0.386. The trend of decreasing density with age is clear, and fit is reasonable (*r*^2^ = 0.6140). The curvature for later ages is driven by the single older sample (Figure 2b, right-most point), and so quantitative results using this regression should be interpreted with care. We believe that this method is conservative and underestimates an age correction, as the density decrease from the younger samples is much stronger than that indicated by the single older sample.

Using the methods above, we estimate that the human age of 45 years old corresponds to a macaque age of 14.8 years old, much older than the mean macaque sample age of 5.2 years old. The ratio of expected callosal density for younger vs. older samples is 1.2, indicating that if a younger human sample was used, we estimate the density would be about 20% higher. This 20% correction drives our statement that human callosal fiber count is likely closer to 240 million fibers, rather than 200 million.

Aboitiz et al. reported an (uncorrected) fiber density in the human corpus callosum of 371.7 fibers/*µ*m^2^ (their Table 1); applying all three corrections (‘371.7 * 0.65 * 1.44’) we obtain a value of 347.9 fibers/*µ*m^2^. To compare to the estimate from our regression, we plug a value of 1350 grams for the average human brain size into the allometric equation and obtain an estimate of 309.1 fibers/*µ*m^2^ for the fiber density of the human corpus callosum. The difference between the two estimates is approximately 11.5%.

To further investigate how the corrected human data point compares with the Wang et al. data, we recomputed the allometric regression of callosal axon density with the Wang et al. data with the corrected human datapoint. The regression equation changes to −0.274 with *r*^2^ = 0.928. This is a small (2.5%) change in exponent, compared to −0.281 in Figure 1, with virtually no change in multiplicative constant (*<* 1%), and an improvement in the *r*^2^ value. See supplemental Figure S2.

## Discussion

We noted a confound of age between human and animal samples, and used macaque monkey data to estimate the magnitude of that effect. We conservatively estimate that human callosum density has been underestimated by 20%, because of this uncorrected age confound. We showed that the human data, with all corrections applied to it, fits with our estimate based on smaller-brained animal data. We also showed that including the human data in the allometric regression procedure does not change the regression in any appreciable way.

From this age correction, we estimate that the human corpus callosum has 240 million fibers in it, or 20% more than previously reported. The widely cited estimate of 200 million fibers in the human corpus callosum is taken from this sample of older humans (27), and so is underestimated due to this age confound.

### Comparing estimates of total intrahemispheric and interhemispheric connectivity

Rilling & Insel estimated grey matter surface area and corpus callosum surface area and made an inference about interhemispheric and intrahemispheric connectivity from the data. At least two things prevented them from estimating actual connectivity from their surface area data using these equations. First, at the time of their publication, data had been published and allometric regressions performed for all quantities above except one: connection density within the corpus callosum. This issue was resolved by the allometric regression performed above, based on high-quality electron microscopic data that appeared after Rilling & Insel published their results.

Though allometric regressions had been published on all other variables, a second issue is that some papers only reported the allometric exponent (B in *y* = *Ax*^*B*^, or *log*(*y*) = *A* + *B * log*(*x*) when transformed into log-log space for the linear regression)–sufficient to examine scaling relationships, but insufficient for quantitative estimates of other values. If all of the data from these publications were publicly available, it would be possible to compute parameters for each allometric regression and plug all quantities into the equations.

Unfortunately, no original data was available from any authors we contacted; instead, we use the same methods outlined above to estimate the data and estimation error from the published graphs. We also use data from tables where possible, and cross-reference and combine results across publications where applicable. From these data, we were able to compute a reduced major axis regression on log-log data for all quantities required by equations 1-3. References for the data used, the results of these regressions, and a detailed account of why each dataset was selected can be found in Table S2.

## Results

Plugging in values from Table S1 into Equations 1 and 2, we compute the total number of callosal fibers to scale as the 0.643 power of the total number of white matter fibers (Figure 3).

**Fig. 3.**
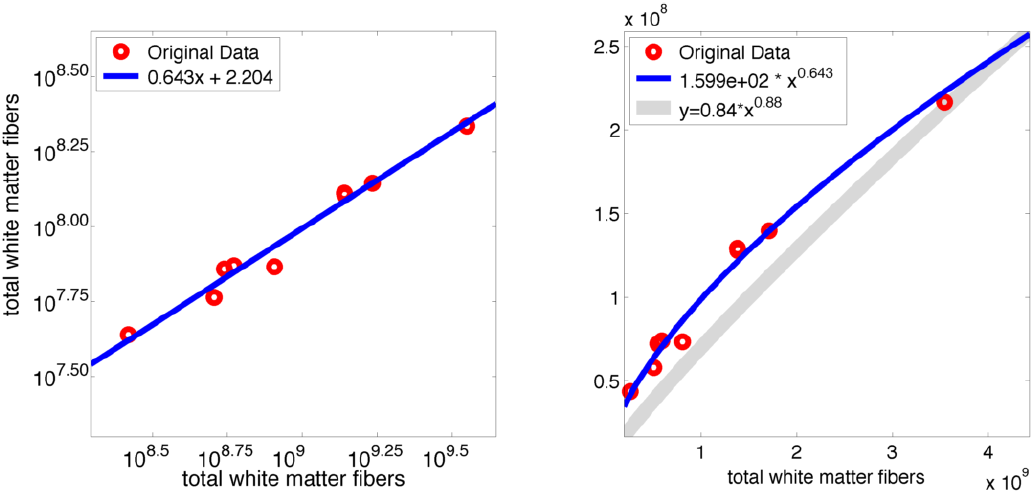
Log-log and euclidean plots of allometric regression for total white matter fibers vs. total callosal fibers. Estimates of connectivity computed from Rilling & Insel’s data are in red, computed allometric regression is in blue. Grey corresponds to the regression equation estimated by Rilling & Insel. The regressions are plotted together to visually show the differences in scaling (curvature); note that the actual values output by their equation are not meaningful in its usage here.

## Discussion

This result (exponent = 0.643) is drastically different than that estimated by Rilling & Insel (1999a) (exponent = 0.88), and suggests that human interhemispheric connectivity is shrinking even faster than they estimated. We suggest that this large deviation from unity indicates that total connectivity may simply be the wrong measure to compare functional connectivity across species. Consistent with this, Shen et al (2014) show no evidence of reduced functional interhemispheric connectivity across primate species, including macaque and human (44).

Indeed, there are no studies that compare total connectivity across species; instead, studies compare between specific fiber bundles (44, 45). These fiber bundles are thought to interconnect specific cortical areas; in fact, a number of cortical parcellation schemes use anatomical and functional connectivity measures to define cortical areas (46, 47). Here we suggest that the relationship between the number of cortical areas and interhemispheric connectivity is a more appropriate measure. Thus, we turned our attention back to the literature to find data that would allow us to compare an inter-area fiber count.

### Relating anatomical and functional connectivity

#### Explaining results through the homotopic connectivity of the corpus callosum

White matter fiber tracts connect as bundles of fibers between two cortical areas. When considering a single cortical area, callosal connectivity is largely to one other cortical area (the homotopic area in the other hemisphere (40)), whereas intrahemispheric connectivity connects many other cortical areas.

Critically, while interhemispheric connections remain homotopic with brain size, the number of intrahemispheric cortical areas that a single cortical area connects to increases with brain size. This is partly driven by an increase in the number of cortical areas in larger brains (48) and is potentially related to “efficient interconnectedness” (49). See Figure 4 for a visual depiction of this.

**Fig. 4.**
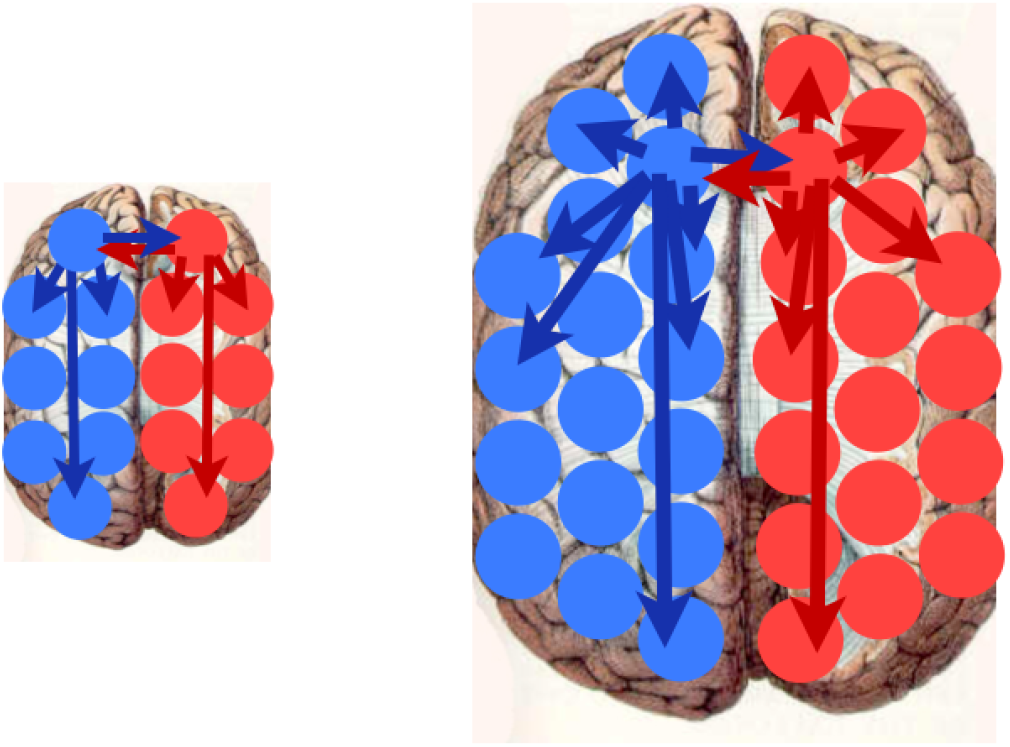
Larger brains have both more cortical areas, as well as more *intrahemispheric* inter-area connections per area. The number of *interhemispheric* inter-area connections per area is the same: one.

This suggests that any individual inter-area connection– whether interhemispheric or intrahemispheric–is a proportionally smaller amount of the total connectivity with increasing brain size, despite the fact that the size of each area, and number of fibers within that inter-area connection is likely increasing with brain size (49, 50).

## Materials & Methods

In order to examine whether this effect can explain the above apparent reduction in interhemispheric connectivity, we directly compared the scaling of fiber connections to the scaling of inter-area connections. For the fiber ratio, we were able to avoid using the estimated scaling equation for the number of cortical areas (48) by computing ratios of per-area quantities. The scaling equation for the number of cortical areas was not well-fit by a power-law equation and the values of its parameters have not been verified in other studies. We also chose to compare callosal connectivity to intrahemispheric connectivity, rather than comparing callosal connectivity to total connectivity. In allometric regression, when regressing quantities A (here callosal connectivity) versus A+B (total connectivity), it has been shown that regressing A versus B (intrahemispheric connectivity) is a more effective method. This approach was used in some (but not all) of the original Rilling & Insel analyses.

In order to compare these scaling laws, we regressed them against each other (as done in Rilling & Insel). In all regressions performed, allometric regression exponents were close to one, and so linear regressions were computed as well to test whether allometric or linear regression was a better fit. However, since computing linear regressions opens up the possibility of non-zero y-offsets (i.e. that the curves do not intersect with the origin), we also estimated best fits for exponential relationships with a y-offset (*y* = *A* x*^*B*^ + *C*) using the Nelder-Mead optimization procedure in MATLAB. Table S3 contains the quantities used in equations 4 and 5 below.

If the relationship between relative fiber count scaling and relative inter-area scaling is linear, this would suggest that the scaling of the number of fibers per intrahemispheric and interhemispheric inter-area connection is the the same. Because an allometric function for fiber counts was estimated above, and an allometric function for per-area inter-area connections is available, it is possible to compute an estimate of the relative fiber counts for callosal vs. intrahemispheric inter-area connections.

Using Equations 1 and 2 and basic algebra, the following equations are obtained:

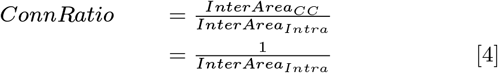

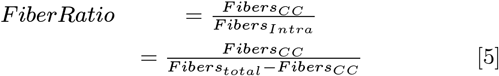

Just as before, by computing the ratio of callosal and intrahemispheric quantities, the allometric equation for the number of cortical areas cancels out. Note that, for this regression only, the percentage of neurons that project into the white matter is critical (see this variable’s value and notes about the value’s selection in Table 1). Variations in that value are noted in the discussion below.

## Results

For the regression of relative fiber scaling and relative inter-area scaling, an equal fit (log-log: *r*^2^ = 0.927; linear: *r*^2^ = 0.928) is obtained with linear regression (assumed exponent = 1) and allometric regression (exponent = 1.207). Though the allometric is not so far from unity, the discrepancy between the allometric and linear exponents leave open the question of whether this relationship is linear or not. The unconstrained optimization (Nelder-Mead method) returns an exponent of 1.039 (*r*^2^ = 0.929) when run on the raw data, lending support that the data have a linear relationship for all relevant species, and that the relationship may not hold (or the data may be noisiest) for the smallest-brained species.

Regressing the expressions in Equations 4 vs. 5 also returns good fits for both allometric (exponent = 1.222, *r*^2^ = 0.977) and linear regressions (*r*^2^ = 0.980). The unconstrained optimization returns an exponent of 1.0343, again consistent with the results from the first regression. The multiplicative value of the linear regression (4.3; see the legend in Figure 5) gives the relative strength of callosal connections vs. intrahemispheric connections.

**Fig. 5.**
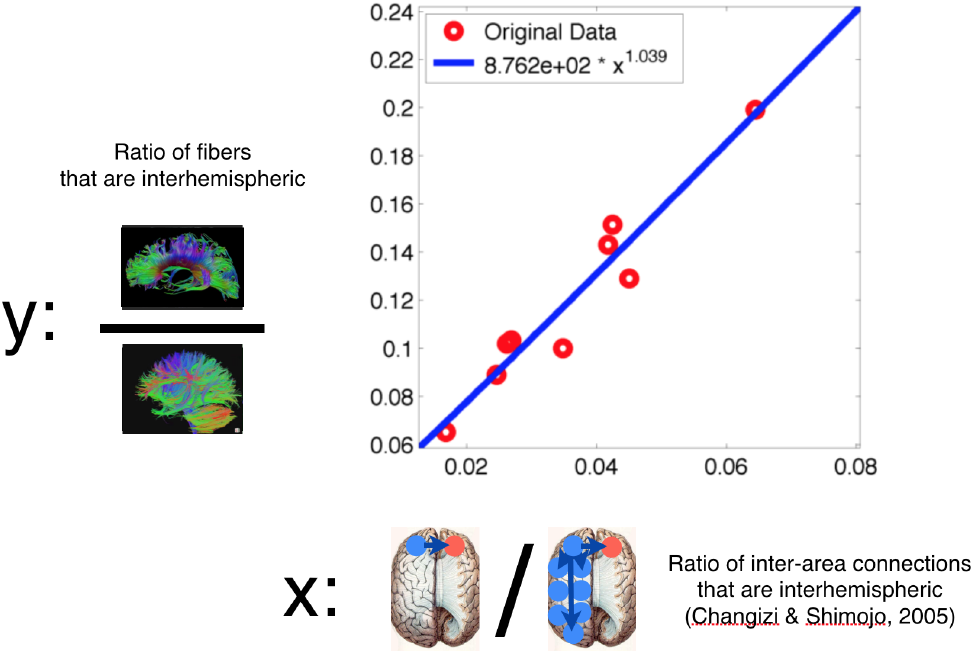
Regressions for callosal fiber count (per inter-area connection) vs. intrahemispheric fiber count (per inter-area connection). The allometric and linear regressions have very similar fits. The multiplicative constant in the linear regressions indicates the relative number of fibers in average callosal vs. intrahemispheric inter-area connections.

What drives the relative reduction of callosal fiber count? As shown above, total callosal connection count scales more slowly than total white matter connection count. The linearity of the first regression here is consistent with the hypothesis that this effect can be explained completely by the differential scaling of callosal vs. intrahemispheric inter-area connections, and that there is no differential scaling of fiber counts within each inter-area connection.

Fiber ratio and inter-area connection ratio showed a linear relationship; there is no exponential component of their relationship. Instead, their relationship is a constant multiplicative factor over brain sizes. This means that the fiber scaling differences found above are completely explained by changes in the relative numbers of inter-area connections. In other words, we expect the proportion of interhemispheric inter-area connections will decrease with brain size there is always only one per cortical area (the connection to the homotopic area in the other hemisphere) while we know that the number of areas, and intrahemispheric inter-area connections are both increasing (48). The exponential rate at which that relationship changes is the same exponential rate at which we computed the number of interhemispheric vs. intrahemispheric fibers to be changing. There is no evidence of selective reduction in the number of fibers per inter-area connection as suggested by Rilling & Insel; increases or decreases in fiber counts exactly correspond to increases or decreases in inter-area connections.

### Relative fiber count of inter- vs intrahemispheric inter-area connections

Comparison of fiber counts, on a per-inter-area connection basis, also showed a linear relationship. This again supports the idea that the same scaling laws apply to both fiber types. Callosal fibers showed 4.3x more fibers per inter-area connection than intrahemispheric connections, suggesting a special role for interhemispheric fibers. This result is even stronger in the light of findings from Henry Kennedy’s lab. They have published data that suggest fiber count decreases as an exponential function of distance (6). Because callosal connections are on average longer than intrahemispheric connections (51), our a priori expectation should be that callosal fiber counts would not be equal, but would be relatively lower when compared to intrahemispheric fiber counts (per inter-area connection).

If we accept this consistent relationship, then the slope of the regression represents the relative number of fibers in a callosal inter-area connection vs. an intrahemispheric inter-area connection. The slope value is 4.3x both in the linear and unconstrained 3-parameter optimization, suggesting that callosal inter-area connections have about 4.3x more connections than the average intrahemispheric inter-area connection. This 4.3x factor is preserved across all brain sizes.

### Potential explanations for a non-linear relationship

The possibility remains that the relationship between fiber and inter-area connection scaling is not a linear relationship. A potential interpretation of the 1.2 exponent value is that it reflects the scaling of non-homotopic connectivity in the corpus callosum. We assumed that all connections are homotopic, but we know that is not strictly true. In addition, it seems likely that there will be a positive scaling of non-homotopic connections with brain size, as the number of non-homotopic areas increases with brain size (48).

It is also worth noting that we used a constant percent of neurons projecting into the white matter, to be conservative. Herculano-Houzel et al. (52) found that the percentage of neurons that project into white matter decreases with brain size, which would increase the positive exponent we’ve found.

### Summary & Conclusions

These analyses suggest that callosal connections are both very typical and very special. There is no evidence for a selective decrease in callosal connections; rather callosal connections scale just like intrahemispheric connections. Across species, callosal connections severely break the exponential distance relationship; despite being on average longer connections, the number of fibers is 4.3x greater in a callosal inter-area connection than in an intrahemispheric inter-area connection.

### General discussion and conclusions

In this paper, we used data extracted from the literature to investigate allometric scaling of total and interhemispheric white matter fibers and inter-area connections across primate species. Along the way, we found that:

- Callosal fiber density scales proportional to brain mass with an exponent of *≈* 0.28 (see Figure 1).
- There is good evidence for age-related decreases of fiber density in macaque monkeys.
- Human callosal axon density is underestimated due to an age confound in the data; we estimated a 20% age-related correction, or that young adult humans have *≈* 240 million callosal fibers.

These findings allowed us to examine cross-species scaling of the proportion of total and interhemispheric white matter fibers. We found that:

- Interhemispheric fibers scale as the 0.64 power to total fibers, much smaller than the 0.88 power previously estimated (2).
- The scaling of interhemispheric vs. intrahemispheric fiber counts appears to match the scaling of interhemispheric vs. intrahemispheric inter-area connections (i.e. fiber tract bundles between areas), suggesting that the above is due purely to the homotopic nature of interhemispheric connectivity.
- When estimating bundle sizes, despite on average being longer connections, callosal fiber bundles contain 4.3x more axons than the average intrahemispheric connection.

### Potential challenges and caveats

One possible challenge to performing the analyses done here is that combining these data mathematically across studies requires challenging cross-study alignment of the data. Choices such as species sampled, tissue preservation and processing, and imaging techniques can all have large effects on individual allometric estimates; because of this, these scaling laws are usually understood more qualitatively than quantitatively. Though we have seen some papers combine data across studies (7, 49, 50, 53), most combine exponents through very rough qualitative means rather than through the quantitative plug-and-play procedures here. We have not seen any quantitative examinations of either approach, though we approach both with caution.

We chose to do this analysis for a number of reasons. First, we believe Rilling & Insel’s interpretation of their data doesn’t account for important scaling differences between the corpus callosum and intrahemispheric inter-area connections, and that when this is accounted for, their data lead to the unlikely (and opposing) hypothesis that fiber connections in callosal inter-area connections are proportionally increasing vs. intrahemispheric inter-area connections. Second, the authors implicitly used similar computational procedures to come to their conclusions. Lastly, we believe our methodology is the best available approach for doing cross-study alignment, has never been presented before, and is therefore a good alternative to existing publications.

As outlined throughout the paper, there are a number of challenges in this type of estimation:

- Without direct access to source data, it is hard to know what data is best to use.
- Allometric regressions are highly-dependent on variables that we don’t have control over when collecting data across the literature.
- There is a lack of convergence in the literature about the allometric relationship for both neural density and the percentage of neurons sending projections into the white matter fiber, both of which affect the 4.3x ratio of intra-vs. inter-hemispheric connectivity.

Allometric and linear regressions equally explained the data with different exponents; which choice reflects the true relationship in the data? The use of the non-parameterized optimization procedure gave weak evidence in favor of the linear regression, and there is theoretical reasoning behind the linear regression, but there is no stronger evidence in support of using either regression at the moment.

### Applicability to other large-brained species

Not only does this work clarify the role of the corpus callosum in humans, but it also re-calibrates expectations for other large-brained animals. With a dissociation between brain size, lateralization, and independence, there is no reason to expect reduced interhemispheric communication, nor lateralization, in large-brained species such as elephants and cetaceans. In animals such as cetaceans where a reduced corpus callosum is clear (54), this should be viewed as unexpected and a specialization. In dolphins, relatively independent hemispheres may allow them maintain a constant state of alertness and ability to surface to breathe, even during their unique sleep pattern where only one hemisphere enters sleep at a time (55).

## Conclusions

We have argued elsewhere that the special conditions of human development lead to a unique developmental trajectory of white matter, particularly the corpus callosum, which affects the development of lateralization (14). We believe the results in this paper, in tandem with our previous results, put a focus back on the human corpus callosum and its role in human lateralization and cognition.

This paper makes a number of testable predictions about the corpus callosum across many species. We orators with access to freshly deceased human, cetacean, or other brains to collaborate with us in testing the hypotheses outlined in this paper.

## ACKNOWLEDGMENTS

This work was partly funded by a Center for Academic Research and Training in Anthropogeny (CARTA) fellowship, as well as by NSF grant SMA 1041755 to the Temporal Dynamics of Learning Center, an NSF Science of Learning Center.

## Footnotes

∗ It is worth noting that the original Rilling & Insel paper, and therefore this one, is focused on white matter connections. This discussion therefore ignores other types of intrahemispheric connectivity: local, inter- and intra-laminar connections and long-distance horizontal projections. Both of these make connections predominantly within a single cortical area and, by definition, never exit the grey matter into the white matter.

† http://github.com/guruucsd/CallosalData

‡ Data from (35) were published after this research was originally completed, and a request for that data has not been returned.

## Notes

### Competing Interest Statement

The authors have declared no competing interest.

